# ONE HEALTH APPROACH ON SARS-COV-2 – USING SHEEP AS SENTINEL ANIMALS TO INCREASE FUTURE PANDEMIC PREPAREDNESS – a pilot study

**DOI:** 10.1101/2024.03.15.585163

**Authors:** Milena Samojlović, João Mesquita, Sérgio Santos-Silva, Malin Neptin, Joakim Esbjörnsson

## Abstract

Coronaviruses are a family of viruses that can infect a number of species of birds and mammals with great zoonotic potential to cross species barriers and cause spill-over events. SARS-CoV-2 has been shown to cause clinical and inapparent disease and mortality in several animals cohabitating with humans. Sheep are also susceptible to SARS-CoV-2 and have potential to harbor and spread the virus, as well as develop neutralising antibodies due to similarities of virus-receptor interactions to those in humans. The main aim of this study was to investigate the prevalence of SARS-CoV-2 neutralising antibodies in sentinel animals after natural exposure to the virus. The serum samples were collected from sheep in Central Portugal, Serra da Estrela region, both prior to and during the COVID-19 pandemic. The sheep were kept on dairy farms for production of Serra da Estrela cheese, in small herds and in constant contact with farm workers. The sera were tested using already established SARS-CoV-2 pseudovirus systems for multiple SARS-CoV-2 variants including Wuhan, Delta and Omicron. Partial neutralisation activity towards Wuhan and Delta variants was observed, while neutralisating antibody escape was observed in all Omicron variants tested due to the mutations present . Our results indicate that potential SARS-CoV-2 virus cross-species transmission could have been established through contacts between people and animals on sheep farms. Using farm animals as sentinels is of great importance for implementing One Health Approach in zoonotic virus surveillance and control towards increasing future pandemic preparedness.

## Introduction

Coronaviruses (CoV) are a family of viruses that can infect a number of species including birds, mammals, humans, and that have the potential to diversify and cross species barriers [1]. The recent SARS-CoV-2 pandemic was caused by a coronavirus of zoonotic origin [2] and that has led to enormous global health problems and challenges, economic hardships and social insecurities over the past four years. The emergence of new SARS-CoV-2 variants due to increased virus fitness and number of mutations led to more effective transmission and immune escape [3]. SARS-CoV-2 has been shown to cause clinical and inapparent disease and mortality in several animals cohabitating with humans. So far, the virus has been reported in 18 animal species from ten families (*Felida, Viverridae, Hyaenidae, Canidae, Mustelidae, Procyonidae, Cervidae, Hippopotamidae, Hominidae, and Cricetidae*) [4,5]. The circulating SARS-CoV-2 variant infections have been reported in animals during pandemic, where Delta variant as the most dominant was detected in 15 animal species, more than one variant was detected in ten animal species and wild type virus was detected in eight animal species [4].

Sentinel animal for SARS-CoV-2 could be any animal in which changes in known characteristics (levels of antibodies) can be measured to assess interrelationships between human and animal health regarding the virus circulation and interspecies transmission and to provide an early warning system of those implications [6]. Wild ruminants as white-tailed deer have been reported to be very susceptible to SARS-CoV-2 infection, being infected with four different variants: Alpha, Delta, Gamma, and Omicron [4, 7, 8]. White-tailed deer is capable of deer-to-deer virus transmission even suggesting possibilities of deer-to human transmission [9]. Recent studies have shown that sheep are susceptible to SARS-CoV-2, both by supporting the virus replication in ovine derived cell cultures and by experimental infection [10, 11]. Seroconversion was also reported in infected sheep [11, 12]. Other studies showed that sheep have potential to harbor and spread the virus, as well as develop neutralising antibodies due to similarities of virus-receptor interactions to those in humans [12, 13].

Sheep farming in Portugal has a long tradition especially in rural areas and therefore the selection has resulted in local native sheep breeds specialized for either meat, milk or wool [14]. One of the most important dairy sheep breeds is Serra da Estrela, inhabiting the Mountain region of the same name. The milk is used for production of famous Portuguese Serra da Estrela cheese with Protected Designation of Origin. The sheep are held extensively or semi-extensively in herds of different sizes with an average herd size consisting of little over 100 animals [15]. With semi-extensive grazing regime and milking performed two times per day by all farmers highly significant contacts between people and sheep and between grazing animals and potential wildlife could have been established on these farms.

Potential SARS-CoV-2 virus cross-species transmission could have occurred through contacts between people and sheep during daily animal handling and milking procedures, as well as between gazing animals and susceptible wildlife, especially during pandemic. The main aim of this study was to investigate the prevalence of anti-SARS-CoV-2 neutralising antibodies in sentinel animals. The blood serum was collected from Serra da Estrela sheep in Portugal both prior to and during the COVID-19 pandemic. More specifically, we aimed to investigate whether different SARS-CoV-2 variants circulating in people at certain time points had different influence in eliciting the neutralising antibody responses in sheep at corresponding time points.

## Material and methods

### Sample collection and study region

Sheep serum samples were collected from the autochthonous sheep breed (Serra da Estrela) from dairy farms in Serra da Estrela Region, Central Portugal, used for production of Serra da Estrela cheese. The sheep were kept in small herds and were in constant contact with farm workers, but no data is available on whether some of the workers had Covid-19 infection during the pandemic. Samling was carried out by official veterinarians for the national brucellosis surveillance program. Blood was collected following the official regulations and protocols ensuring animal welfare. In the laboratory blood serum was separated after centrifugation (2500xg, 5 min), aliquoted and stored at -20°C until testing. A total of 180 serum samples were collected at three different time periods. Prepandemic samples (45 samples) were collected in 2016 long before SARS-CoV-2 pandemic was introduced and were investigated for the presence of neutralising antibodies against Wuhan variant, the earliest SARS-CoV-2 variant. Pandemic samples (58 samples) were collected in January 2022 and were investigated for the presence of neutralising antibodies towards Delta (30 samples) and Omicron BA.1 variants (28 samples), circulating just before and during sampling period, respectively. Lastly, pandemic samples collected in March 2023 (77 samples) during the circulation of XBB Omicron variant were investigated against it (47 samples) and Omicron BQ.1.1 variant (30 samples) circulating just before the sampling period.

### Pseudovirus production and pseudovirus neutralisation assay

We investigated the presence of neutralising antibodies against SARS-CoV-2 virus in sheep serum samples from Portugal using SARS-CoV-2 pseudovirus systems for multiple SARS-CoV-2 variants. Briefly, pseudovirus constructs (PV) have been developed by three-plasmid co-transfection in HEK293 cells using lentiviral backbone (Addgene # 8455), firefly luciferase reporter (Addgene #170674) and S protein expressing plasmids of different SARS-CoV-2 variants - Wuhan, Delta, Omicron BA.1, XBB, BQ.1.1(Addgene #170442, #172320, #180375, #194494, #194493) by Lipofectamine 3000 as a transfection reagent [16, 17, 18, 19, 20]. Pseudovirus particles were collected after 60 hours from supernatant, filtered, aliquoted and frozen at -80 °C until use. The vesicular stomatitis virus (VSV) positive pseudovirus control was constructed similarly by replacing the S protein expressing plasmid with G envelope protein expressing plasmid pCMV-VSV-G.

HEK293 and HEK293-ACE2 cells were cultured in Dulbecco’s Modified Eagle Medium (DMEM) with high glucose and sodium pyruvate, 10% fetal calf serum and 100 IU/mL penicillin and 100 μg/mL streptomycin. Cells were propagated in incubator at 37°C with 5% CO2.

The pseudovirus neutralisation assay was performed in flat bottom 96-well plates in HEK293-ACE2 cells by previously described protocol [21]. All serum samples were inactivated at 56°C for 1 hour prior to testing. Three-fold serially diluted sheep sera samples were tested in duplicates and incubated for 1 hour at 37°C with SARS-CoV-2 pseudoviruses sufficient to produce approximately 100000 relative light units in 50µl. Negative and positive controls were also included in the testing. After initial incubation and addition of 20000 HEK293-ACE2 cells per well, plates were incubated for 72 hours at 37°C. Luminescence was measured using Bright-Glo Luciferase Assay System (Promega, Madison, WI, USA) in black 96-well plates using VICTOR® Nivo™ Plate Reader (PerkinElmer, USA).

### Statistica Analysis

Neutralisation was quantified as the reduction in luciferase activity relative to the average of six control wells infected with pseudoviruses in the absence of serum and plotted against the logarithm of the dilution factors using GraphPad Prism software (San Diego, CA, USA). A four-parameter logistic equation was fitted to the neutralisation curves.

## Results and discussion

Forty-five prepandemic samples were tested for the presence of neutralising antibodies towards earliest SARS-CoV-2 Wuhan variant. Pandemic samples collected during January 2022 were tested towards Delta variant and Omicron BA.1 variant, 30 samples and 28 samples respectively. Pandemic samples collected in March 2023 were tested towards Omicron variants, 47 and 30 samples towards XBB and BQ.1.1 variants respectively. The pandemic sheep serum samples were tested against officially reported variants circulating in human population just before or during the period of sampling. Adjusting the SARS-CoV-2 pseudovirus system used for neutralising antibody screening to the virus variants circulating just before or during sampling periods decreases the possibility of false-negative results due to SARS-CoV-2 antibody escape in new variants. Moreover, making SARS-CoV-2 pseudovirus system variant-specific lowers the possibility of cross-reactivity to other non-targeted coronaviruses because of the presence of specific mutations in SARS-CoV-2 variants, especially Omicron that influence antibody response [22]. The frequency of SARS-CoV-2 variants in human population in Central Portugal during SARS-CoV-2 pandemic is shown in Figure 1 (data collected by the Portuguese National Institute of Health available on their official webpage <u>SARS-CoV-2 Portugal (insa.pt)</u>).

**Figure 1.**
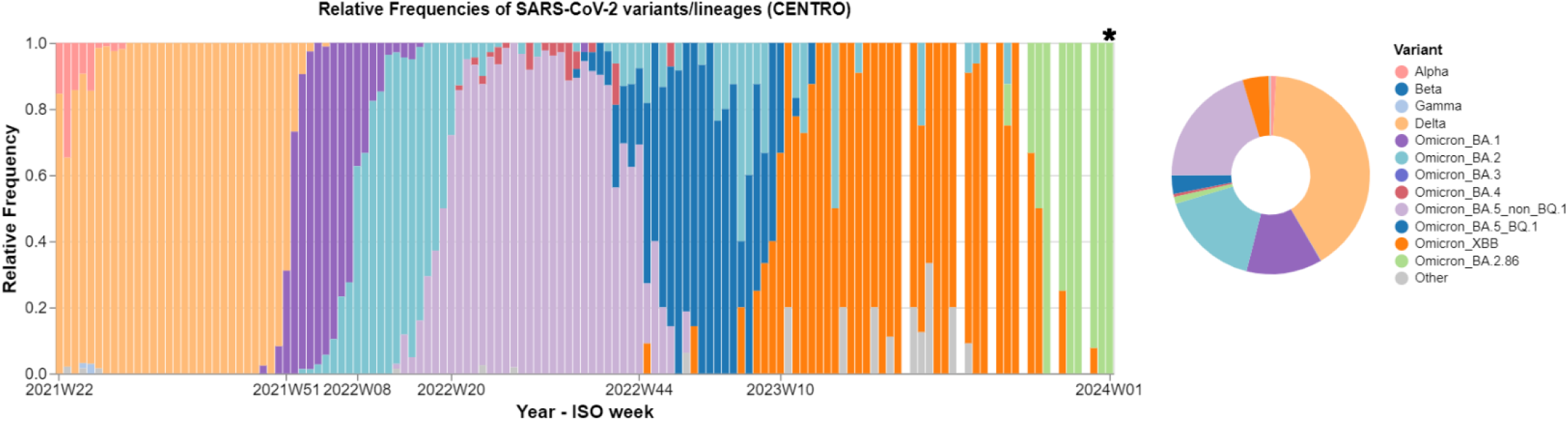
Overview of weekly relative frequency of SARS-CoV-2 variants during 2021-2023 in Central Portugal. In the period of first pandemic sampling in January 2022 (Week 1 to week 4) the most dominant SARS-CoV-2 variant circulating was Omicron BA.1 preceded by Delta variant. In the period of second pandemic sampling in March 2023 (week 9 to week 13) the most dominant variant circulating was Omicron XBB preceded by Omicron BA.5_BQ1. (source: <u>SARS-CoV-2 Portugal (insa.pt)</u>, Accessed 28^th^ December 2023)

The results of neutralisation capacities of tested sheep sera towards SARS-CoV-2 are presented in Figure 2. The sheep sera were tested with an initial dilution of 1:3 followed by 3-fold serial dilutions. The differences in the neutralisation response curves can be observed between prepandemic and pandemic samples, as well as between different SARS-CoV-2 variants both with sheep sera samples and WHO anti-SARS-CoV-2 antibody standards. As expected, WHO standard for anti-SARS-CoV-2 immunoglobulin for early variants (21/340) showed complete neutralisation activity against Wuhan variant PV, while WHO standard for anti-SARS-CoV-2 immunoglobulin for Omicron variant (21/338) showed complete neutralisation against Omicron BA.1 PV while antibody escape was detected against Omicron XBB PV. None of the sheep serum samples tested showed 100% neutralization against SARS-CoV-2 variants in any of the dilutions. In prepanedmic samples only one sample showed 81% neutralisation against Wuhan PV in lowest (1:3) dilution, while other six samples showed neutralization between 60-70% in the same dilution with neutralisation percentage gradually decreasing as dilution increased. Non-specific background of animal sera in SARS-CoV-2 pseudovirus based neutralization assay in lower dilutions was reported by other authors where they detected around 70% neutralisation in lowest dilution in mouse SARS-CoV-2 negative sera. Same authors report around 60% neutralisation in lowest dilution in human negative sera as well [23]. Moreover, in prepandemic healthy donor sera tested against SARS-CoV-2 by cell- and virus-free S protein-based neutralisation assay, neutralisation of around 60% to SARS-CoV-2 was seen in initial dilutions as well [24]. In our study higher background of prepandemic sheep sera could have been also due to the potential previous exposure of sheep to bovine coronavirus which was shown to cross-react with neutralisation antibodies to SARS-CoV-2 [25]. In pandemic samples partial neutralisation activity towards Delta variant was observed, whereas neutralisating antibody escape was observed for all the tested Omicron variants. Delta variant antibody respionse is more similar to the one that was elicited by early 2020 variant such as Wuhan [26]. SARS-CoV-2 antibodies escaped to Omicron variants were previously described in human sera samples [27, 28]. Highest number of samples (7) with >80% neutralisation in lowest dilution was dectected in 2022 panedemic samples towards Delta variant PV in comparison to samples tested to Wuhan and Omicron variant PV. This was also the SARS-CoV-2 variant tested with the lowest number of samples (5) that showed neutralisation of <50% in initial dilutions. Pandemic samples from 2022 tested for the presence of neutralising antibodies towards Omicron BA.1 variant showed 5 samples with >70% neutralisation in lowest dilution, while more than half the samples showed neutralisation below 50% in all dilution tested. Pandemic samples from 2023 showed highest number of samples below 50% neutralisation in all dilution, 36 of 47 samples tested against XBB variant and 29 of 30 samples tested against BQ.1.1. Only two samples showed partial neutralisation of 82% and 72% in lowest dilution towards XBB and BQ.1.1, respectively.

**Figure 2.**
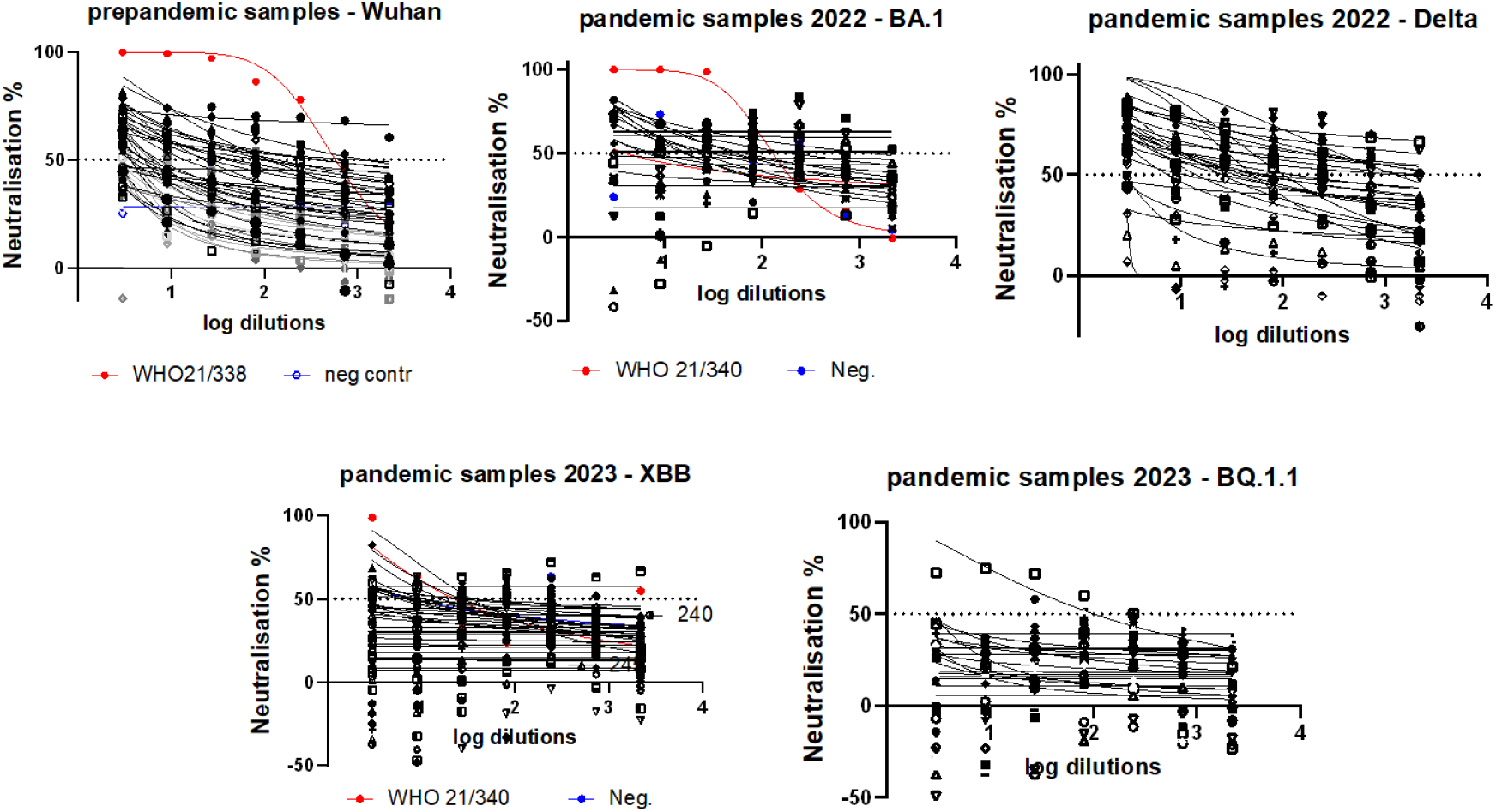
SARS-CoV-2 Pseudovirus neutralisation assay results of sheep sera samples in HEK293-ACE2 cells. Neutralisation capacities of sheep sera sampled prepandemic and during pandemic in January 2022 and March 2023 towards SARS-CoV-2 variants circulating at sampling time (black lines) and WHO International Standards for anti-SARS-CoV-2 immunoglobulin for early(21/340) and Omicron (21/338) variants (red lines).

Sheep can be experimentally infected with SARS-CoV-2 and as a result of the infection, serological response was deteced by both ELISA and VNT. More seropositive sheep after experimental SARS-CoV-2 infection was detected by ELISA, while only one sheep had low levels of neutralising antibodies of 1:20 titer value [11]. Presence of neutralising antibodies in sheep serum and even colostrum and milk after immunization with fusions of the SARS-CoV-2 receptor binding domain (RBD) to ovine IgG2a Fc domains were reported by Jacobson et al, 2023 [29]. However, the experimantal exposure of animals to virus is not always comparable to the exposure in natural conditions due to different evolution and pathogenesis of natural infection, especially when it comes to the virus dose the animals were exposed to, which is always higher in experimantal conditions [30]. This further emphasize the importance of field studies and serological monitoring of virus circulation in natural conditions.

In our study only partial neutralisation of SARS-CoV-2 varinats was seen by sheep serum samples after natural exposure to SARS-CoV-2. The seroprevalence of SARS-CoV-2 in sheep naturally exposed to the virus was examined by other authors by different serological method. Villaneuva-Saz et al, 2021 reported no positive samples in sheep that were in close contacts with veterinary student community in Spain during pandemic by In-House ELISA targeting IgG specific for receptor biding domain region of S protein [30]. The first serological evidence of SARS-CoV-2 natural infection in sheep and goats in Italy was reported by Fusco et al, 2023 by commercialy available ELISA and virus neutralisation test (VNT). By ELISA, they detected more positive sheep in comparison to goats, with only one sheep being positive by VNT with neutralisation titer value at the limit of detection (1:20) [12]. To our knowledge no sheep serum samples were tested towrds SARS-CoV-2 before our study by PV neutralisation test available for different SARS-CoV-2 variants. Eventhough the results have shown that potential SARS-CoV-2 virus cross-species transmission could have been established through contacts between people and sheep during pandemic, further serological analysis needs to be undertaken.

The study emphasizes the need for addressing some of One-Health challenges like disease outbreak and surveillance through the development of effective and efficient translational diagnostic tools for zoonotic viruses, and through establishing an interdisciplinary scientific collaboration. An important part of pandemic preparedness is to create more rapid responses between pathogen emergence and development of diagnostics and medical countermeasures, and implementing research outcomes effectively to both human and animal population, especially in the fight against zoonotic viruses with pandemic potential.

## Conclusion

The fast mutations rates of SARS-CoV-2 and occurrence of new virus variants have been shown to have influenced its transmissibility to different animal species as well. Using animals cohabitating in close contact with humans as sentinels represents great potential for more comprehensive zoonotic virus surveillence for identifying potential high-risk areas, and informing evidence-based strategies for disease prevention and control. Especially we would like to point out the advantage of using farm animals for zoonotic virus surveillance since serum samples can be easily accessible due to the national animal diseases monitoring and surveillance programs requested by the government or collected at the slaughterhouses before animals being processed for meat production. No specific ethical permission is required, and the serum volume is usually sufficient to perform several tests. Implementation of One Health Approach is of crucial importance for the increased and better future pandemic preparedness.

## Author Contributions

Conceptualization, M.S., J.E. and J.M.; methodology, J.E. and M.S.; validation, J.E. and M.S.; formal analysis, M.S.; investigation, M.S.; resources, J.E., M.S, J.M., S.S.-S., and M.N.; writing—original draft preparation, M.S.; writing—review and editing, J.E., J.M., M.S. and S.S.-S.; visualization, M.S.; supervision, J.E.; project administration, M.S., J.E. and M.N.; funding acquisition, M.S. and J.E. All authors have read and agreed to the published version of the manuscript.

